# Transition State Interactions in a Promiscuous Enzyme: Sulfate and Phosphate Monoester Hydrolysis by *Pseudomonas aeruginosa* Arylsulfatase

**DOI:** 10.1101/327643

**Authors:** Bert van Loo, Ryan Berry, Usa Boonyuen, Mark F. Mohamed, Marko Golicnik, Alvan C. Hengge, Florian Hollfelder

**Affiliations:** Department of Biochemistry, University of Cambridge, Cambridge, United Kingdom; Department of Chemistry and Biochemistry, Utah State University, Logan, Utah 84322, United States

## Abstract

*Pseudomonas aeruginosa* arylsulfatase (PAS) hydrolyses sulfate and, promiscuously, phosphate monoesters. Enzyme-catalyzed sulfate transfer is crucial to a wide variety of biological processes, but detailed studies of the mechanistic contributions to its catalysis are lacking. We present an investigation based on linear free energy relationships (LFERs) and kinetic isotope effects (KIEs) of PAS and active site mutants that suggest a key role for leaving group (LG) stabilization. In LFERs wild type PAS has a much less negative Br0nsted coefficient (β_leaving group_^obs-Enz^ = −0.33) than the uncatalyzed reaction (β_leavingroup_^obs^ = −1.81). This situation is diminished when cationic active site groups are exchanged for alanine. The considerable degree of bond breaking during the TS is evidenced by an ^18^O_bridge_ KIE of 1.0088. LFER and KIE data for several active site mutants point to leaving group stabilization by active-site lysine K375, in cooperation with histidine H211. ^15^N KIEs combined with an increased sensitivity to leaving group ability of the sulfatase activity in neat D_2_O (Δβ_leaving group_^H-D^ = +0.06) suggest that the mechanism for S-O_bridge_ bond fission shifts, with decreasing leaving group ability, from charge compensation via Lewis acid interactions towards direct proton donation. ^18^O_nonbridge_ KIEs indicate that the TS for PAS-catalyzed sulfate monoester hydrolysis has a significantly more associative character compared to the uncatalyzed reaction, while PAS-catalyzed phosphate monoester hydrolysis does not show this shift. This difference in enzyme-catalyzed TSs appears to be the major factor favoring specificity toward sulfate over phosphate in this promiscuous hydrolase, since other features are either too similar (uncatalyzed TS) or inherently favor phosphate (charge).

## INTRODUCTION

Arylsulfatases catalyze the *in vivo* hydrolysis of sulfate monoesters, producing inorganic sulfate, typically removing it from a sugar or a steroid hormone. Sulfatases are highly proficient enzymes, with catalytic proficiencies ((*k*_cat_/*K*_M_)/*k*_uncat_) well above 10^13^-10^17^ M^−1^ for the model substrate 4-nitrophenyl sulfate **1d** (Scheme 1).^1–3^ Despite their occurrence in eukaryotes and prokaryotes, relevance for a variety of key processes (e.g. development,^4–10^ germination,^11^ resistance against toxic defense molecules,^12,13^ mucin desulfation,^14–16^ or degradation of mucopolysaccharides^17,18^) and the occurrence of various diseases as a result of their malfunction (e.g. lysosomal disorders^17,19^), their mechanism has not been studied in the same detail as that of the related phosphatases.

**Scheme 1.**
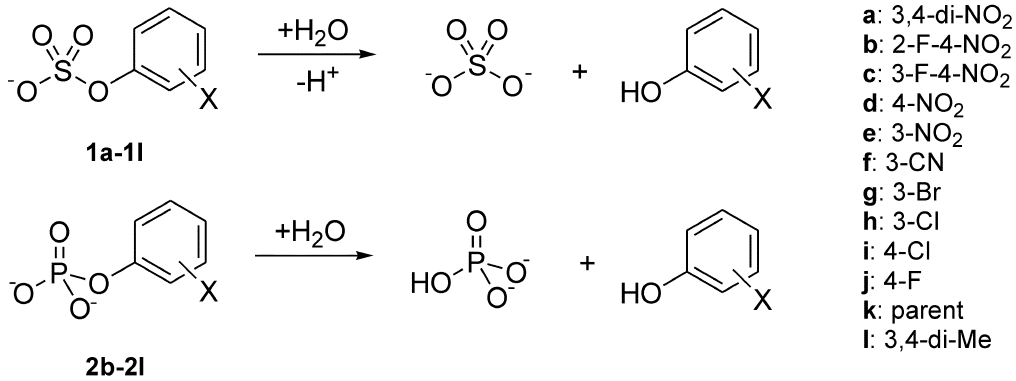
General reaction scheme for the PAS-catalyzed hydrolysis of aryl sulfate monoesters **1a**-**1l** and aryl phosphate monoesters **2b**-**2l**.

The majority of sulfatases known to date are members of the alkaline phosphatase (AP) superfamily. The mechanism of transition state (TS) stabilization during enzyme-catalyzed substrate hydrolysis of one member of this superfamily, *Escherichia coli* alkaline phosphatase (*Ec*AP), has been subject of a large number of in-depth studies involving the experimental tools of linear free energy relationships (LFERs), kinetic isotope effects (KIEs), mutant studies, and structural analysis by X-ray crystallography.^20–36^ In addition computational simulations have been employed to pin-point transition state interactions.^26,27,37–41^

Sulfatase members of the AP superfamily have been studied less extensively. Catalytically important residues have been shown to be conserved among the arylsulfatases^2,42–47^ and mutant studies (e.g. of human arylsulfatase A,^48^ choline sulfatase,^47^ and the closely related phosphonate monoester hydrolase^49^) have suggested that many of these conserved residues are indeed involved in the catalytic pathway in the enzyme active site. LFERs and KIEs have been measured for a number of phosphatases,^20–23,35,50–58^ but only one such study is available for sulfatases.^59^

In addition to the family relationship, members of the AP superfamily are also typically catalytically promiscuous,^60–62^ i.e. they catalyze multiple, chemically distinct reactions with large rate accelerations.^63,64^ Within the superfamily reciprocal relationships of crosswise catalytic promiscuity are observed, i.e. the promiscuous activity of one member is the native function of another, and *vice versa*.^63,64^ Given the postulated role of promiscuity in evolution by gene duplication,^60,65^ functional crossover defines such functional relationships as pathways for respecialization or repurposing in enzyme superfamilies.^66^ As catalysis for the activity under selective pressure must be maintained at a relevant level during evolution, the question of how mechanistic features of these enzymes can be effective for catalysis of different reactions arises.

We have studied the reactions catalyzed by the promiscuous arylsulfatase from *Pseudomonas aeruginosa* (PAS) using LFERs and KIEs. PAS has a wide active site opening^42^ and - in contrast to sulfatases with high specificity for a particular leaving group (such as a sugar moiety^67,68^ or choline^47^) - accepts a range of aromatic substrates: so that the construction of LFERs based on series of aryl sulfates of different reactivity is possible with minimal interference from unique binding effects. PAS operates at high rates (*k*_cat_/*K*_M_ 4.9×10^7^ s^−1^M^−1^), even for the hydrolysis of the non-natural substrate 4-nitrophenyl sulfate **1d.**^1^ In addition, PAS promiscuously catalyzes the hydrolyses of phosphate mono-,^1^ and diesters^62^ as well as phosphonates,^69^ thus covering mechanistically distinct hydrolase reactions.^70^ However, despite its catalytic promiscuity, PAS is a genuine sulfatase: it is typically expressed under sulfate starvation conditions, part of an operon coding for sulfate-processing enzymes, and thought to act as a sulfate scavenger.^71,72^

Based on the X-ray structure of PAS^42^ (and those of the human arylsulfatases A^45,73^ (hASA) and B^43^ (hASB) combined with mutant data for hASA^48^) a double displacement catalytic pathway was proposed, in which a post-translationally modified cysteine,^74^ formylglycine fGly51, performs a nucleophilic attack on the sulfur center (Figure 1b). The covalent hemiacetal intermediate is broken down with assistance of H115 acting as a general base (step 2 in Figure 1b). Finally the aldehyde form of the fGly nucleophile is again hydrated, regenerating the enzyme for the next round of catalysis. Mutant data for several of the analogous active site residues in hASA show that a single mutation into an alanine of one of the residues likely to be involved in charge compensation during the TS results in lowered but still detectable levels of activity.^48^ During the hydrolysis of sulfates, negative charge is expected to build up in the TS on the sulfuryl group and on the leaving group.^3,75^ TS stabilization can be achieved by offsetting this charge build-up: using positively charged functionalities such as metal ions, or by using hydrogen bonding or electrostatic interaction with positively charged amino acid side chains. The latter possibly may involve proton transfer to the leaving group oxygen. The active site of PAS contains a number of residues that could neutralize charge build-up on oxygen atoms during the transition state, or affect the p*K*_*a*_ of the formylglycine (fGly51) nucleophile (Figure 1).

**Figure 1.**
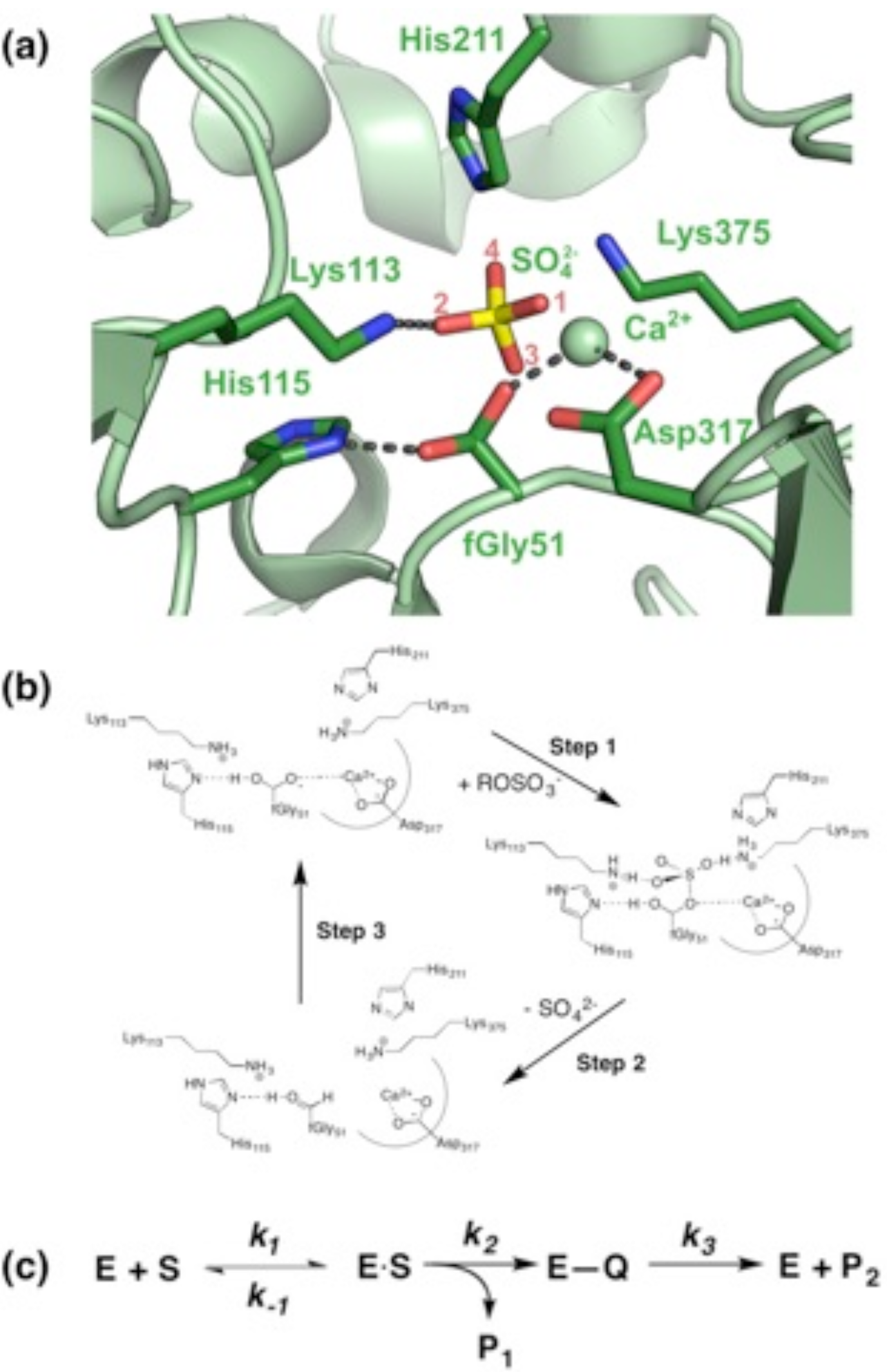
Active site and reaction mechanism of PAS. (**a**) 3D representation of the active site of PAS with a bound sulfate ion.^42^. The assignment of the analogous non-bridging (O_1_-O_3_) and bridging (O_4_) oxygens in the sulfate ester substrate were based on positions of the active site residues in the X-ray structure of human arylsulfatase A C69A^73^ with *p*-nitrocatechol sulfate bound in the active site. (**b**) Proposed catalytic pathway for PAS-catalyzed sulfate monoester hydrolysis. Upon binding of the sulfate monoester substrate, the hydrated fGly51 performs a nucleophilic attack on the sulfur atom and the bond to the alcohol leaving group (ROH) is broken (S-O_bridge_ bond fission) (Step 1). The covalent intermediate is broken down by base-catalyzed hemiacetal cleavage in which inorganic sulfate acts as the leaving group (Step 2). The enzyme is subsequently regenerated by hydration of the formylglycine aldehyde (step 3). (**c**) Schematic representation of the steps in panel **b**. *k*_1_ is the rate of formation of the enzyme-substrate (E.S) complex from free enzyme (E) and substrate (S) and *k*_−1_ represents the dissociation rate of the ES-complex. *k*_2_ is the rate constant for the nucleophilic attack of the hydrated formylglycine and subsequent S-O_bridge_ bond fission (step 1), and *k*_3_ that of hemiacetal cleavage (step 2). The rehydration of the fGly residue (step 3) is expected to be several orders faster that hemiacetal cleavage, and thus was not considered for the interpretation of pre-steady state kinetics. Product P_1_ is the phenolate leaving group expelled from the substrate in step 1; product P_2_ is inorganic sulfate.

Here we provide a detailed quantitative examination of their influence on the nature of the TS in the arylsulfatase-catalyzed reaction. We use LFER and KIE data to compare the nature of the TS for the native PAS-catalyzed sulfate monoester hydrolysis with the uncatalyzed reaction, and show that, while both are dissociative, the TS of the enzyme-catalyzed reaction has a more associative character. The latter difference is absent for phosphate monoester hydrolysis. Pre-steady-state kinetic data provide information about the rate-limiting step in the overall kinetic mechanism. Finally, changes in LFERs and KIEs as a result of alanine scanning mutagenesis of PAS active site residues suggest that Lys375 serves as the general acid that minimizes leaving group charge change in the TS and provides leaving group stabilization.

## RESULTS AND DISCUSSION

### Steady state parameters

In order to obtain insight into the nature of the transition state (TS) of PAS-catalyzed sulfate monoester hydrolysis we used a series of substrates with varying leaving groups to measure linear free energy relationships (LFERs), similar to those constructed for several phosphoryl transfer enzymes^21,22,31,32,55–58,70,76^.

Purified PAS WT^1^ was used to determine Michaelis-Menten parameters *k*_cat_, *K*_M_ and *k*_cat_/*K*_M_ for a series of phenyl sulfate monoesters (compounds **1a-1l**, Scheme 1) with varying leaving group abilities (represented by their p*K*_a_ values, which range from 5.2 to 10.35 (Table S1 and Figures S1 and S2, supporting information, SI)). The steady-state catalytic rate constant (*k*_cat_) is independent of leaving group ability (Figure 2a, varying between 9-18 s^−1^; Table S1, SI). Based on the kinetic scheme for enzyme-catalyzed substrate hydrolysis (Figure 1c), the relevant rate constant to probe for information on the nature of the TS should be a good proxy for *k*_2_. If enzyme-catalyzed cleavage of the sulfate ester bond were fully rate-limiting, the steady state catalytic rate constant *k*_cat_ would be essentially equal to *k*_2_ (equation S14, SI). If this is not the case, i.e. if *k*_2_ ≥ *k*_3_, more complex terms arise (see equations S1-3, SI). Assuming that the substrate binding constants (*K*_D_ = *k*_−1_/*k*_1_) are similar for the complete series of sulfate monoesters (Scheme 1), *k*_cat_/*K*_M_ reports on all steps from free enzyme and substrate to the first irreversible transition state. In this case the formation of the covalent intermediate between the formylglycine nucleophile and the respective sulfur center of the substrates that leads to leaving group departure is likely to be irreversible, as no significant inhibition by the phenolate product is observed. (See equations S1-S3 in the SI for details on the relation between the Michaelis-Menten parameters and the individual rate constants for the various enzymatic steps).

**Figure 2.**
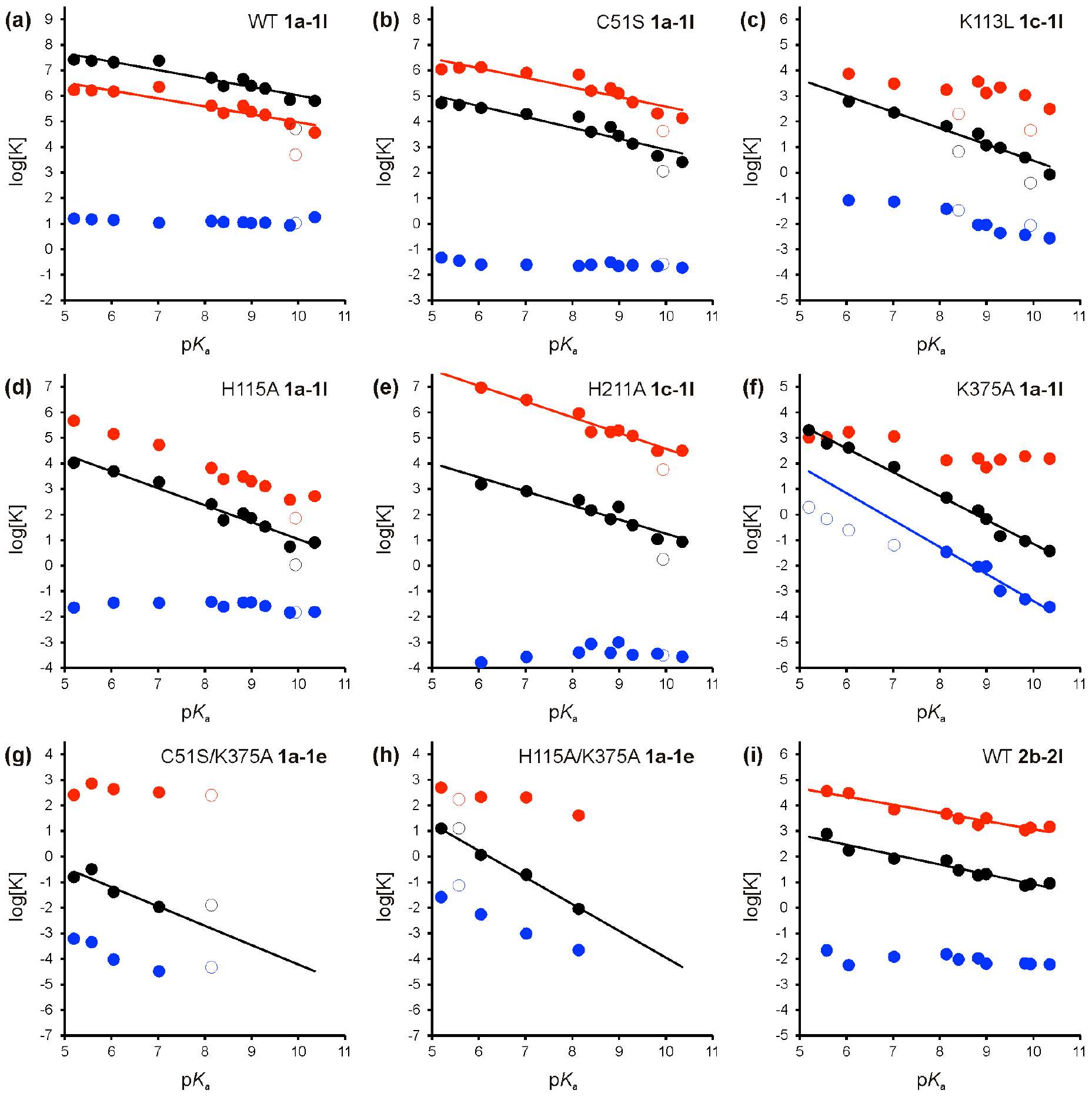
Dependence of catalytic parameters *K*_cat_ (blue) 1/*K*_M_ (red) and *K*_cat_/*K*_M_ (black) (log-values) for PAS-catalyzed hydrolysis of sulfate monoesters on leaving group ability (as represented by the p*K*_a_ of the free phenol in solution). (**a**) PAS WT; (**b**) PAS C51S; (**c**) PAS K113L; (**d**) PAS H115A; (**e**) PAS H211A; (**f**) PAS K375A; (**g**) PAS C51S/K375A; (**h**) PAS H115A/K375A. (**i**) PAS WT with phosphate monoesters. All data were obtained in 100 mM Tris-HCl pH 8.0; 500 mM NaCl at 25 °C. The resulting slopes (=β_leaving group_^obs^) are listed in Table 1 and S14. The data for sulfate monoester **1k** deviated significantly from the trend for all enzymes, suggesting a consistent difference in binding constant as compared to the other sulfate monoester substrates and were therefore not included in any of the fits. Data points represented by open circles were not included in the fits. All data for the kinetic parameters are listed in the supporting information (Table S1, S3-S10).

**Table 1.** Overview of the observed Brønsted constants for leaving group dependence (β_leaving group_^obs^) of *K*_cat_/*K*_M_ for PAS-catalyzed sulfate and phosphate monoester hydrolysis.^a^

| Reaction | catalyst | $\beta_{\text{leaving group}}^{\text{obs}}$ |
| --- | --- | --- |
| Sulfate monoesters | Solution (neutral) | -0.27<sup>b</sup> |
|  | Solution (monoanion) | -1.81<sup>b</sup> |
|  | WT | -0.33±0.04 |
|  | C51S | -0.43±0.05 |
|  | K113L | -0.64±0.06 |
|  | H115A | -0.67±0.05 |
|  | H211A | -0.55±0.07 |
|  | K375A | -0.94±0.04 |
|  | C51S/K375A | -0.76±0.24 |
|  | K113L/K375A | nd<sup>c</sup> |
|  | H115A/K375A | -1.04±0.06 |
|  | H211A/K375A | nd<sup>c</sup> |
| Phosphate monoesters | Solution (monoanion) | -0.27<sup>d</sup> |
|  | Solution (dianion) | -1.23<sup>d</sup> |
|  | WT | -0.39±0.04 |

<sup>a</sup>All enzymatic reactions performed in 100 mM Tris-HCl pH 8.0, 500 mM NaCl, 25 °C. <sup>b</sup>Edwards *et al.*<sup>3</sup>, H<sub>2</sub>O as the nucleophile. <sup>c</sup>Not determined. Activity too low to be detectable. <sup>d</sup>Kirby & Vargolis<sup>86</sup>, H<sub>2</sub>O as the nucleophile.

### Attempts to determine pre-steady state kinetics

Direct determination of pre-steady state kinetic parameters such as *k*_2_ using stopped-flow methods requires a detectable burst phase. When the PAS WT-catalyzed hydrolysis of sulfate monoester **1d** was monitored at millisecond time-scale using a stopped-flow setup, only linear time courses without a burst phase were observed. This scenario does not rule out the existence of a burst, since a large ratio *k*_2_/*k*_3_ would make the relevant burst phase undetectable. When the time axis was corrected for the dead time of the equipment (3 ms), extrapolation of the linear time courses did not converge to the same point at t = 0 (Figure S3a, SI). The values for the positive intercepts at the product axis were directly correlated with the amount of active enzyme that was added (Figure S3b, SI), indicating the existence of a burst phase lasting less than 3 ms. This observation confirms an overall rate-limiting step that occurs after sulfate ester bond cleavage. Simulations of the stopped-flow time courses based on values for *K*_D_, *k*_2_ and *k*_3_ that fit the steady state data (equations S5-7, Figure S4, SI) suggested that the actual *K*_D_ for substrate binding of sulfate monoester **1d** is at least 100 μM (i.e. >150-fold higher than its *K*_M_). Simulation of the LFERs for *k*_cat_ and 1/*K*_M_ based on the same three pre-steady state parameters confirmed this observation (assuming *K*_D_ is equal for all substrates, Figure S5, SI). Consistent with *k*_3_ being rate-limiting for the entire LFER, the apparent *K*_M_ values for the substrate series increased with increasing leaving group p*K*_a_ (from 0.6-28 μM, Table S1, SI) and log plots of *k*_cat_/*K*_M_ and 1/*K*_M_ versus leaving group p*K*_a_ showed similar slopes of −0.33±0.04 and −0.31±0.04 respectively (Figure 2a). Since *k*_cat_ is more or less equal to *k*_3_, which is independent of intrinsic substrate reactivity, it cannot be used as a proxy for the rate relevant to the LFER. We therefore use *k*_cat_/*K*_M_ as a proxy for *k*_2_.

### LFER for PAS WT-catalyzed sulfate monoester hydrolysis

Plots of *k*_cat_/*k*_M_ against the leaving group p*k*_*a*_ (Figure 2) were linear, with a β_leaving group_^obs^ of −0.33 that was considerably less negative than that of the rate for the uncatalyzed hydrolysis reaction (*k*_uncat_), for which a β_leaving group_^obs^ of −1.81 has been measured.^3^ Williams *et al.*^59^ recently constructed a LFER for PAS, but arrived at a much steeper slope (−0.86). Although most points superimpose well with our data (see Figure S6, SI), the choice of bulky leaving groups with higher p*k*_*a*_ (specifically the inclusion of the bulky 4-amino-acetyl- and 4-methoxyphenolate), the narrower range of p*k*_a_-values (three compared to almost five log units covered here) as well as basing the study on overall fewer data points (7 *vs* 12 in our work) with worse significance (p-value, 0.026 *vs*. <10^−4^) and correlation coefficients (R^2^, 0.66 *vs*. 0.91) seem to have resulted in a distortion of the slope due to idiosyncratic effects of substrates (e.g. due to steric clashes). However, in both cases the slope β_leaving group_^obs^ is less steep than that for the solution reaction.

A possible cause for the considerably less negative β_leaving group_^obs^ compared to the solution reaction could be that the *k*_cat_/*K*_M_-values do not only represent a chemical step, as shown previously for wild-type *E. coli* alkaline phosphatase (AP).^34,52,77^ Fast enzymatic reactions (*k*_cat_/*K*_M_ ~10^6^-10^9^ s^−1^ M^−1^) can be diffusion controlled, i. e. a physical step that occurs prior to the first chemical step can be rate-limiting. The *k*_cat_/*K*_M_-values for the PAS WT-catalyzed hydrolysis of sulfate monoesters **1a**-**1l** ranged from 10^5^-10^7^ s^−1^ M^−1^ (Table S1), which partially fall in the range mentioned above. In terms of the physical and chemical steps that occur during the reaction cycle of PAS (Figure 1c), diffusion control of *k*_cat_/*K*_M_ arises when *k*_1_×[S] is smaller than *k*_2_ (which means *k*_−1_ ≪ *k*_2_). In extreme cases *k*_cat_, *K*_M_ and *k*_cat_/*K*_M_ can even be completely independent of *k*_2_ (equations S11-13, SI). Since *k*_2_ is expected to decrease with increasing p*k*_a_, this extreme scenario is most likely for substrates bearing leaving groups with low p*k*_a_, resulting in a flattening of the LFER. In this case the resulting β_leaving group_^obs^ will be less steep than the actual β_leaving group_ of the chemical reaction of interest (step 2 in Figure 1b). Diffusion controlled reactions are known to be slowed down when performed in increasingly viscous conditions. Viscosity dependence data for the PAS WT-catalyzed conversion of sulfate monoesters **1c** and **1d** showed a decrease in *k*_cat_/*K*_M_ with increasing sucrose concentration (Figure S7). The decrease was the result of a decrease in *k*_cat_, whereas *K*_M_ was unchanged. However, based on equations S1-3 (SI) *K*_M_ is expected to increase, whereas *k*_cat_ is expected to be unaffected by changes in the rate of substrate binding (*k*_1_×[S]) or dissociation of the enzyme-substrate (ES) complex (*k*_−1_), which are the steps expected to be affected by increased viscosity. Slowing down the dissociation of the final sulfate product would lower *k*_3_ and thereby *k*_cat_. However, *K*_M_ would decrease in this scenario as well (according to equation S9) and *k*_cat_/*K*_M_, which is independent from *k*_3_, would be unchanged: yet this is not observed (Figure S7). At the high sucrose concentration used here it is not unlikely that even a weak molecular binding event causes inhibition. The data could be fitted to an inhibition constant of ~1.1 M for both *k*_cat_ and *k*_cat_/*K*_M_ for both substrates. This suggests pure non-competitive inhibition of PAS by sucrose (20% (w/v) = 0.58 M), which would explain the observed lowering of *k*_cat_ and *k*_cat_/*K*_M_ while *K*_M_ is unaffected (see the legend to Figure S7, SI, for more details). To test whether the LFER for PAS WT-catalyzed sulfate monoester hydrolysis was affected, data were recorded in 10% (w/v) sucrose and showed a β_leaving group_^obs^ identical to the one performed in the absence of the viscogen (Figure S8). Since the viscosity dependence is strongest for faster reactions, a diffusion-controlled reaction is expected to show a *less* negative β_leaving group_^obs^ in media with increased viscosity. We therefore conclude that the observed kinetic parameters that is relevant for monitoring leaving group effects for PAS WT-catalyzed hydrolysis of sulfate monoesters **1a**-**1l** are not diffusion controlled and that the deviation of β_leaving group_^obs^ from the β_leaving group_ of the solution reaction is a genuine effect of substrate binding and turnover by the enzyme.

Previous experimental studies into the nature of the TS of enzyme-catalyzed sulfate transfer using LFERs^59,78^ typically show less steep correlations than the corresponding solution reaction,^79–82^ bringing the Brønsted slopes to values closer to zero. This decrease could be ascribed to interactions with cationic groups in the active site, but no KIEs or mutational data were available to check this hypothesis. In addition a LFER for sulfate transfer has been reported for the promiscuous sulfatase activity of AP WT.^30^ Subsequent KIE studies showed that in this case a dissociative TS is likely.^20^ In AP the less negative β_leaving group_^obs^ compared to the solution reaction was explained by interaction of the leaving group with positively charged moieties in the enzyme active site, most likely a divalent metal ion (Zn^2+^). The latter phenomenon has also been reported for AP-catalyzed phosphate monoester hydrolysis.^83^ In several protein tyrosine phosphatases a protonated aspartate was identified as responsible for leaving group stabilization.^50,51,53–55,58^ Replacement of this aspartate with asparagine restored the leaving group dependence to a value close to that of the solution reaction.^55^ In protein phosphatase 1 (PP1), a histidine was thought to provide the same role, although practical limitations (low yield and poor activity) prevented experimental verification.^57^

### LFER for PAS WT-catalyzed phosphate monoester hydrolysis

The fact that PAS WT is also a proficient phosphatase^1^ ((*k*_cat_/*K*_M_)/*k*_uncat_ = 2.9×10^11^ M^−1^ towards phosphate monoester **2d**), opens up the possibility to study two reactions that proceed through similar TSs in solution^30,52,75,84–86^ in a single active site, and also facilitates comparisons with the more widely studied phosphatases.

Michaelis-Menten parameters were determined for a series of phosphate monoesters (**2b**-**2l**, Scheme 1). As for the sulfatase reaction, *k*_cat_ is practically independent of the leaving group p*k*_a_ (varying between 0.6-1.2×10^−2^ s^−1^, Figure 2i and Table S3, SI), suggesting that the rate-limiting step for phosphate as well as sulfate monoesters is not leaving group-dependent. The *K*_M_ values increase with leaving group p*k*_a_ and range from 0.03-0.92 mM, around ~100-fold higher than for sulfatase activity. The slope of the Brønsted plot β_leaving group_^obs^ for *k*_cat_/*K*_M_ for phosphate monoester hydrolysis is −0.39±0.04, identical within the error margins to the value observed for the sulfatase reaction (and confirmed by a crosscorrelation graph with a slope of unity, see Figure S9, SI). As observed for the sulfatase reaction, enzyme-catalyzed phosphate monoester hydrolysis is considerably less sensitive to leaving group ability than the solution reaction for the phosphate monoester dianion (β_leaving group_ = −1.23^86^). As for sulfate monoester hydrolysis, these considerable deviations can be caused by stabilization/masking of locally developing negative charge, in particular on the leaving group oxygen. Kinetic isotope effect studies suggested that the TS is similar to the solution reaction (see below for details), as in previously studied phosphatases.^22,36,55,57,58^

### The effect of mutations on the β_leaving group_^obs^ for sulfate monoester hydrolysis

As discussed above, the β_leaving group_^obs^ of enzyme-catalyzed phosphate and sulfate transfer reactions can be influenced by compensation by positively charged moieties in the active site of the negative charge build-up that occurs during the TS.^22,32,36,57,78^ Furthermore the change in nucleophile between the solution (H_2_O) and enzyme-catalyzed (formylglycine) reaction can also influence the β_leaving group_^obs^. Zalatan *et al.*^36^ formulated these considerations in order to be able to calculate expected differences in leaving group dependence between enzyme-catalyzed (β_leaving group_^Enz^) and solution reactions (β_leaving group_^solution^) (equation 1). The expected change in leaving group dependence resulting from a change in nucleophile between the enzyme-catalyzed (fGly) and solution (H_2_O) reaction is calculated from the difference in nucleophilicity (p*K*_nuc_^Enz^ − p*K*_nuc_^solution^) weighed with the sensitivity of the leaving group dependence of the reaction type to a change in nucleophile (*p*_xy_). The second of the contributing factors is leaving group dependent binding of the ground state (GS, β_bind_^GS^) and TS (β_bind_^TS^). For *k*_cat_/*K*_M_-based β_leaving group_ values the latter two are indistinguishable from each other and are treated as a single variable (∑β_bind_= β_bind_^GS^ +β_bind_^TS^).

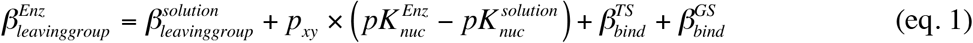

For *E. coli* AP in particular the interaction of the leaving group oxygen with one of the two Zn^2+^ ions during the GS and TS was thought to be mainly responsible for the ∑β_bind_ of +0.33^83^ (3-times larger than the expected contribution of the change in nucleophile from H_2_O to the active site serine of +0.11). Selective removal of only the Zn^2+^-ion responsible for leaving group stabilization is not feasible and therefore experimental assessment of these calculations was not possible. The assignment of the analogous positions of the nonbridging and bridging (leaving group) oxygens in an inorganic sulfate molecule with the active site residues of PAS (as shown in the X-ray structure,^42^ Figure 1), was based on homology with structural data for the enzyme-substrate complex of an inactive variant of human arylsulfatase A.^73^ The majority of the interactions with the nonbridging and leaving group oxygens are expected to be provided by amino acid side-chains, which opens up the possibility of assessing the importance of the correction factors of equation 1 experimentally by determining the leaving group dependence of active site mutants of PAS. To this end the nucleophile (fGly51) and a residue that directly interacts with it (H115) were mutated to assess the contribution of the nucleophile. Furthermore, several positively charged groups expected to provide charge compensation during the GS and TS by interacting with nonbridging (K113 and K375) and leaving group (H211 and K375) oxygens were removed by mutating the respective residues into alanine or leucine.

All mutants were purified to homogeneity and Michaelis-Menten parameters determined for the same series of sulfate monoesters (**1a**-**1l**, Scheme 1) as used with the wild-type enzyme. The mutations resulted in 10^3^-10^8^-fold drops in catalytic efficiencies (*k*_cat_/*K*_M_) for the various substrates (Tables S4-S8, SI). For mutants C51S and H211A *k*_cat_ was still independent of leaving group ability (Figure 2b and 2e respectively), suggesting that, as in the wild-type enzyme, the leaving group-dependent step is not rate limiting. However the *k*_cat_ for these mutants was reduced ~10^3^-fold (C51S) and ~10^5^-fold (H211A) compared to PAS WT. The *K*_M_ values for PAS C51S and H211A were within an order of magnitude of those for the wild-type enzyme (Tables S4 and S7) and, as observed for wild-type PAS, the β_leaving group_^obs^ values for *k*_cat_/*K*_M_ and 1/*K*_M_ are nearly identical. For PAS K113L and H115A both *k*_cat_ and 1/*K*_M_ decreased with increasing leaving group ability (Figure 2c and d, Table S5 and S6), albeit it only at the higher end of the p*k*_a_ spectrum for H115A. For PAS K375A *k*_cat_ decreases with increasing leaving group p*K*_a_, indicating that the rate-limiting step is largely leaving group dependent. The *K*_M_-values are increased ~10^3^-fold compared to the wild-type enzyme and are largely constant (varying in a range of 5-10 mM) for the substrates with a leaving group p*K*_a_ > 8 (Figure 2f, Table S8). The β_leaving group_^obs^ for *k*_cat_/*K*_M_ is nearly identical to that for *k*_cat_ for p*K*_a_ > 8. As for PAS WT, the LFERs for *k*_cat_ and 1/*K*_M_ for PAS K113L, H115A and K375A could be simulated based on assumed values for the pre-steady state kinetic parameters (Figure S10, SI). In particular for K113L and K375A the break in the LFER for *k*_cat_ could be explained by a change of the rate-limiting step with increasing p*K*_a_^leaving group^.

The β_leaving group_^obs^ of all active-site mutants was less negative than the β_leaving group_^obs^ for the wild type. Δβ_leaving group_ (=β_leaving group_^obs-WT^ − β_leaving group_^obs-mutant^) was calculated as the slope of the linear correlation of log[(*k*_cat_/*K*_M_)_WT_/(*k*_cat_/*K*_M_)_mutont_] *vs*. p*K*_a_^leaving group^ (Table 2, Figure S11a). As described above the difference in leaving group dependencies between enzyme-catalyzed and uncatalyzed sulfate and phosphate monoester hydrolysis is influenced by the nature of the nucleophile and charge compensation effects (equation 1). Based on the pH-rate profile for PAS-catalyzed sulfate monoester hydrolysis the p*K*_nuc_ of the enzyme is expected to be < 7.2^1^. The *p*_xy_-value for sulfate transfer is not known, but is expected to be similar to the value for phosphate monoesters (0.013^87^). Assuming that the p*K*_nuc_^Enz^ ~ 6 and p*K*_nuc_^solution^ = −1.7 (nucleophile = H_2_O), the effect of the change in nucleophile between the PAS WT-catalyzed and the solution reaction is expected to be 0.013×(6-(−1.7)) = +0.10. Assuming the difference in nucleophilicity between fGly (solution p*K*_a_ ~ 13-14^88^) and serine (p*K*_a_ ~ 16, similar to that of ethanol) in solution translates into the same difference in the enzyme active site, a small increase in the contribution of the nucleophile term in equation 1 is expected for mutant C51S. Based on this assumption the Δβ_leaving group_^WT-C51S^ is expected to be small and negative. However, we measure a value of +0.10 (Table 2). Since fGly is interacting directly with the Ca^2+^-ion, changing it to a serine may cause a change in the charge compensation effects provided by the divalent cation, which could explain the positive value for Δβ_leaving group_^WT-C51S^. Removal of H115 is expected to result in a slight increase in the p*K*_a_ of the nucleophile, again predicting a small negative Δβ_leaving group_^WT-H115A^ as a result. However, the measured value of +0.32 suggests that any small effect of the mutation on the nucleophile term is overshadowed by a considerable decrease in ∑β_bind_.

The removal of the residues that directly interact with the leaving group oxygen is expected to have a large effect on ∑β_bind_, whereas groups that interact only with the nonbridging oxygens are expected to contribute to ∑β_bind_ at ~10% of the value expected for direct interactions with the leaving group oxygen.^36^ Mutation of H211 resulted in a Δβ_leaving group_^WT-H211A^ of only +0.16, even though this residue interacts exclusively with be the leaving group oxygen. Removal of the nearby K375 results in a Δβ_leaving group_^WT-K375A^ of +0.61. Since the removal of a positive charge is expected to increase the p*K*_nuc_^Enz^, the actual effects on ∑β_bind_ may be slightly higher than observed. These data suggest that K375 is largely responsible for leaving group stabilization. However, unlike in PTPase^55^, where a protonated aspartate has been ascribed to performing the same function, its removal does not result in a near-complete abolition of the difference between β_leaving group_^obs-Enz^ and β_leaving group_^obs-solution^, since Δβ_leaving group_^PAS K375A-solution^ = +0.87. Values of ~+0.1-0.2 would be expected in case of complete removal of charge compensation on the leaving group as a result of the mutation (i.e. ∑β_bind_ = 0, only the change in nucleophile results in a small positive Δβ_leaving group_^obs^). This observation could be rationalized by partial compensation of the loss of charge offset provided by K375 by nearby H211. The same phenomenon (i.e. that K375 partially assumes the role of H211) could explain the relatively small effect of mutation H211A on β_leaving group_^obs^. The combined effects of these two residues find support in the double mutant enzyme PAS H211A/K375A, for which no sulfatase activity was detectable (*k*_cat_/*K*_M_ < 5×10^−6^ s^−1^ M^−1^ for sulfate monoester **1d**, see SI for consideration of the experimental detection limit). This observation strengthens the idea that the effect of each mutation is buffered by the nearby presence of a residue that can take over its function: if the measured reductions in activity for the single mutants were simply additive, the expected *k*_cat_/*K*_M_ for sulfate monoester **1d** should be 2.6×10^−3^ s^−1^ M^−1^ (according to equation S21, SI), and still be detectable. By contrast, their cooperativity leads to a larger detrimental effect, when both are removed.

**Table 2.** Effect of mutations on leaving group dependence for PAS-catalyzed sulfate monoester hydrolysis in PAS WT and K375A

| mutation | reaction | effect in | $\Delta\beta_{\text{leaving group}}^{\text{obs}}$ |
| --- | --- | --- | --- |
| C51S | **1a-1l** | WT | +0.10±0.04<sup>a</sup> |
|  | **1a-1d** | K375A | +0.01±0.33<sup>b</sup> |
| K113L | **1c-1l** | WT | +0.24±0.05<sup>a</sup> |
| H115A | **1a, 1c-1l** | WT | +0.32±0.03<sup>a</sup> |
|  | **1a-1l** | K375A | +0.15±0.05<sup>b</sup> |
| H211A | **1c-1l** | WT | +0.16±0.07<sup>a</sup> |
| K375A | **1a-1e** | WT | +0.61±0.03<sup>a</sup> |

<sup>a</sup>Determined as the slope for $\log[(k_{\text{cat}}/K_{\text{M}})_{\text{WT}}/(k_{\text{cat}}/K_{\text{M}})_{\text{mutant}}]$ plotted vs. leaving group $\text{p}K_{\text{a}}$ (Figure S10a). <sup>b</sup>Determined as the slope for $\log[(k_{\text{cat}}/K_{\text{M}})_{\text{K375A}}/(k_{\text{cat}}/K_{\text{M}})_{\text{double mutant}}]$ plotted vs. leaving group $\text{p}K_{\text{a}}$ (Figure S10b, SI).

As discussed above the effect of leaving group dependent ground and transition-state binding (∑β_bind_) is expected to be modest for residues that interact mainly with the nonbridging oxygens: ~10% (for all the non-bridging oxygens combined) of the value expected for leaving group oxygen^83^. This consideration suggests that the maximal combined effect of K113, K375 and Ca^2+^ on ∑β_bind_ *via* interactions with the non-bridging oxygens is expected to be ~+0.13 (10/110×(β_leaving group_^obs-WT^ − β_leaving group_^obs-solution^) = 0.09×(−0.33-(−1.81))=+0.13). Since removal of a positive charge is expected to increase the p*K*_nuc_ of the enzymatic reaction, the observed effect on ∑β_bind_ of the interaction with non-bridging oxygens, as a result of removing any of these three functional groups is expected to be <+0.05. However, mutation K113L results in a Δβ_leaving group_^WT-K113L^ of +0.24. This large effect partly explains why removal of H115 has an unexpectedly large Δβ_leaving group_^WT-H115A^ of +0.32, since H115 and K113 interact closely in the 3D structure of PAS. The large effect of mutations C51S, K113L and H115A on PAS activity suggests that these mutations influence the interaction of the H211A/K375A pair with the leaving group oxygen. If the effects were electrostatic and isolated from the interactions with the leaving group oxygens, the effect of these mutations would be expected to be identical in K375A and WT, i.e. these three mutations should be additive to mutation K375A. The combined effects of K113L and K375A result in completely inactive enzyme (*k*_cat_/*K*_M_ < 5×10^−6^ s^−1^ M^−1^ for sulfate monoester **1d**), which is lower than the expected value (7.0×10^−4^ s^−1^ M^−1^; based on eq. S21, SI) for an additive effect of both mutations. This can be explained by interaction of both K113 and K375 with the non-bridging oxygens, in which case their simultaneous removal most likely results in complete abolition of substrate binding. The introduction of mutations C51S and H115A into the K375A variant does result in enzymes with detectable activities. However, the effect on the β_leaving group_^obs^ is much lower in the K375A mutant than in the wild-type enzyme, if present at all (C51S in K375A has no significant effect on β_leaving group_, see Table 2 and Figure S10b for details). The non-additive behavior suggests that the unexpectedly large value of Δβ_leaving group_^WT-mutant^ for both these mutations is due to their effect on the interactions between the leaving group oxygen and K375, i.e. this effect cannot be achieved unless K375 is present. Possible explanations for this phenomenon are (*i*) changes in the overall electrostatic character of the active site as result of the mutations that decrease the strength of the interaction between K375 and the leaving group oxygen or (*ii*) changes in substrate positioning that cause the optimal configuration of the K375-leaving group oxygen paring to be distorted, resulting in a lower contribution of this interaction to TS stabilization.

### Kinetic Isotope Effects (KIEs)

Kinetic isotope effects were measured for the bridging (or leaving group) and non-bridging oxygens as well as the nitro group for enzyme-catalyzed hydrolysis of sulfate monoester **1d** and phosphate monoester **2d** (Figure 3; Chart S1, SI), to complement the Brønsted analysis above ^55,57^. The KIE experiment requires turnover of approximately half of a 100 μmol sample of labeled substrate. With mutants of low activity this can require long reaction times, or unpractically large amounts of enzyme. For that reason, most but not all of the mutants for which kinetic data are presented have accompanying KIE data in Table 3.

**Figure 3.**
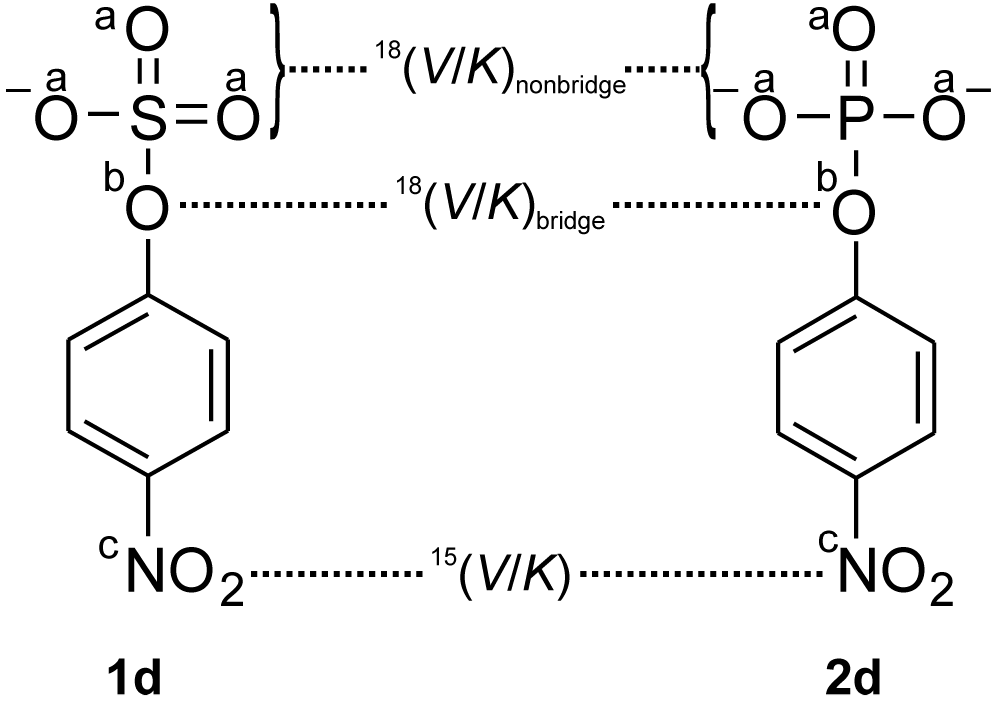
Positions for the kinetic isotope effect measurements in sulfate monoester **1d** and phosphate monoester **2d**. Effects were measured for the nonbridging oxygens (a, ^18^O_nonbridge_ KIE), bridging oxygen (b, ^18^O_bridge_ KIE), and for the nitro group (c, ^15^N KIE). See Chart S1 (SI) for the structures of the labeled compounds used.

**Table 3.** Kinetic isotope effects for PAS-catalyzed hydrolysis of sulfate monoester **1d** and phosphate monoester **2d**.

| substrate | catalyst | $^{15}(V/K)$ | $^{18}(V/K)_{\text{bridge}}$ | $^{18}(V/K)_{\text{nonbridge}}$ |
| --- | --- | --- | --- | --- |
| **1d** | solution (neutral)<sup>a</sup> | 1.0004 (1) | 1.0101 (2) | 1.0098 (3) |
|  | solution (monoanion)<sup>b</sup> | 1.0026 (1) | 1.0210 (10) | 0.9951 (3) |
|  | WT | 1.0006 (4) | 1.0088 (9) | 1.0064 (15) |
|  | C51S | 1.0012 (6) | 1.0097 (19) | 1.0039 (4) |
|  | H115A | 1.0013 (2) | 1.0216 (17) | 1.0004 (6) |
|  | H211A<sup>c</sup> | - | - | - |
|  | K375A | 1.0023 (5) | 1.0244 (23) | 0.9958 (11) |
| **2d** | solution (monoanion)<sup>d</sup> | 1.0004 (2) | 1.0087 (3) | 1.0184 (5) |
|  | solution (dianion)<sup>d</sup> | 1.0028 (2) | 1.0189 (5) | 0.9994 (5) |
|  | WT | 1.0007 (4) | 1.0102 (9) | 0.9912 (8) |

<sup>a</sup>Recorded in 10 N HCl, 15 °C.<sup>92</sup> <sup>b</sup>Recorded at pH 9.0, 85 °C.<sup>92</sup> <sup>c</sup>Activity too low to be determined. <sup>d</sup>Recorded at 95 °C.<sup>52</sup>

Because the KIEs were measured by the internal competition method they describe effects on *k*_cat_/*K*_M_ (or *V*/*K*),^89^ which, as described above, is representative of the rate constant for the first chemical step (*k*_2_ in Figure 1c). Thus, the KIEs determined here reflect the TS for the substrate reacting with the formyl glycine nucleophile, even though the overall rate-determining step is breakdown of the intermediate.*

The expected ranges of the isotope effects in sulfate monoester **1d** and phosphate monoester **2d** and their interpretation have been discussed in detail previously.^90,91^ The secondary KIE at the nitrogen atom, ^15^(*V*/*K*), reports on negative charge development on the nitrophenolate leaving group in the transition state. The *p*-nitrophenolate anion has contributions from a quinonoid resonance form, with decreased N-O bond order and increased N-C bond order. Because N-O bonds are tighter in terms of vibrational frequencies, the nitrogen atom is more tightly bonded in neutral *p*-nitrophenol (or in the *p*NPP substrate) than in the phenolate anion. Thus, the ^15^K EIE for deprotonation of *p*-nitrophenol is normal.(Original reference for this: Hengge, A. C.; Cleland, W. W. Direct measurement of transition-state bond cleavage in hydrolysis of phosphate esters of p-nitrophenol. *J. Am. Chem. Soc*. 1990, *112*, 7421-7422. When protonation or other interactions maintain the leaving group in a neutral state, there is no isotope effect (KIE = unity). This KIE reaches its maximum value of about 1.003, reflecting a full negative charge, when the leaving group in the TS has a very high degree of bond fission and no interactions are neutralizing the resulting charge. The KIE at the bridge oxygen atom, ^18^(*V*/*K*)_bridge_, is a primary isotope effect that arises from S-O or P-O bond fission and is also affected by O-H bond formation, if the leaving group is simultaneously protonated in the TS. Bond fission produces normal isotope effects, primarily due to reduction of the stretching vibration in the TS as the force constant is lowered. Protonation of this atom in the TS gives rise to inverse effects, from the new vibrational modes introduced from the forming bond. A large body of data from phosphate and sulfate ester hydrolysis shows the isotope effect from P-O or S-O bond fission is normally larger in magnitude than the inverse effect from protonation. A normal ^18^(*V*/*K*)_bridge_ effect near its maximum of 1.03 reflects a largely broken S-O or P-O bond in the TS, arising from loss of vibrations involving this bond. In native enzymes utilizing general acid catalysis, or uncatalyzed reactions under acidic conditions, the observed ^18^(*V*/*K*)_bridge_ is reduced by protonation, as shown in Table 3.

The leaving group KIEs ^15^(*V*/*K*) and ^18^(*V*/*K*)_bridge_ report on how leaving group stabilization might be compromised by mutation. Loss of general or Lewis acid catalysis will result in increases in both of these KIEs relative to the native enzyme. The isotope effect on the nonbridging oxygen atoms, ^18^(*V*/*K*)_nonbridge_, monitors the hybridization state of the transferring sulfuryl or phosphoryl group, which affects the P-O or S-O nonbridging bond orders and hence their vibrational frequencies. A loose transition state gives rise to slightly inverse effects as these bond orders increase. This isotope effect becomes increasingly normal (i.e. approaching or exceeding a value of 1) as the transition state grows more associative in nature.

The small ^15^(*V*/*K*) of 1.0006 (Table 3), for PAS WT-catalyzed hydrolysis of sulfate monoester **1d** suggests nearly complete neutralization of the negative charge developing on the leaving group from S-O bond fission. Similar neutralization occurs by intramolecular pre-equilibrium proton transfer during the uncatalyzed hydrolysis of neutral sulfate monoesters under acidic conditions^92^ (^15^(*V*/*K*) = 1.0004). The ^18^(*V*/*K*)_bridge_ KIE for the PAS-catalyzed reaction is similar to that of the uncatalyzed hydrolysis of the neutral monoester^92^ and the AP-catalyzed sulfate monoester hydrolysis^20^ (Table 3, Figure 4a). In both cases significant masking of leaving group charge development occurs due to intramolecular proton transfer, or interaction with a Lewis acid, respectively. The magnitude of ^18^(*V*/*K*)_bridge_ is consistent with significant S-O_bridge_ bond fission concomitant with leaving group stabilization, either by interaction with a positively charged group (Lewis acid) or protonation. A similar observation has been made for phosphate monoester hydrolysis catalyzed by AP,^20^ protein phosphatase 1^57^ (PP-1) and several protein-tyrosine phosphatases (PTPs)^50,51,53,54,93^ (Figure 4a, Table S15).

**Figure 4.**
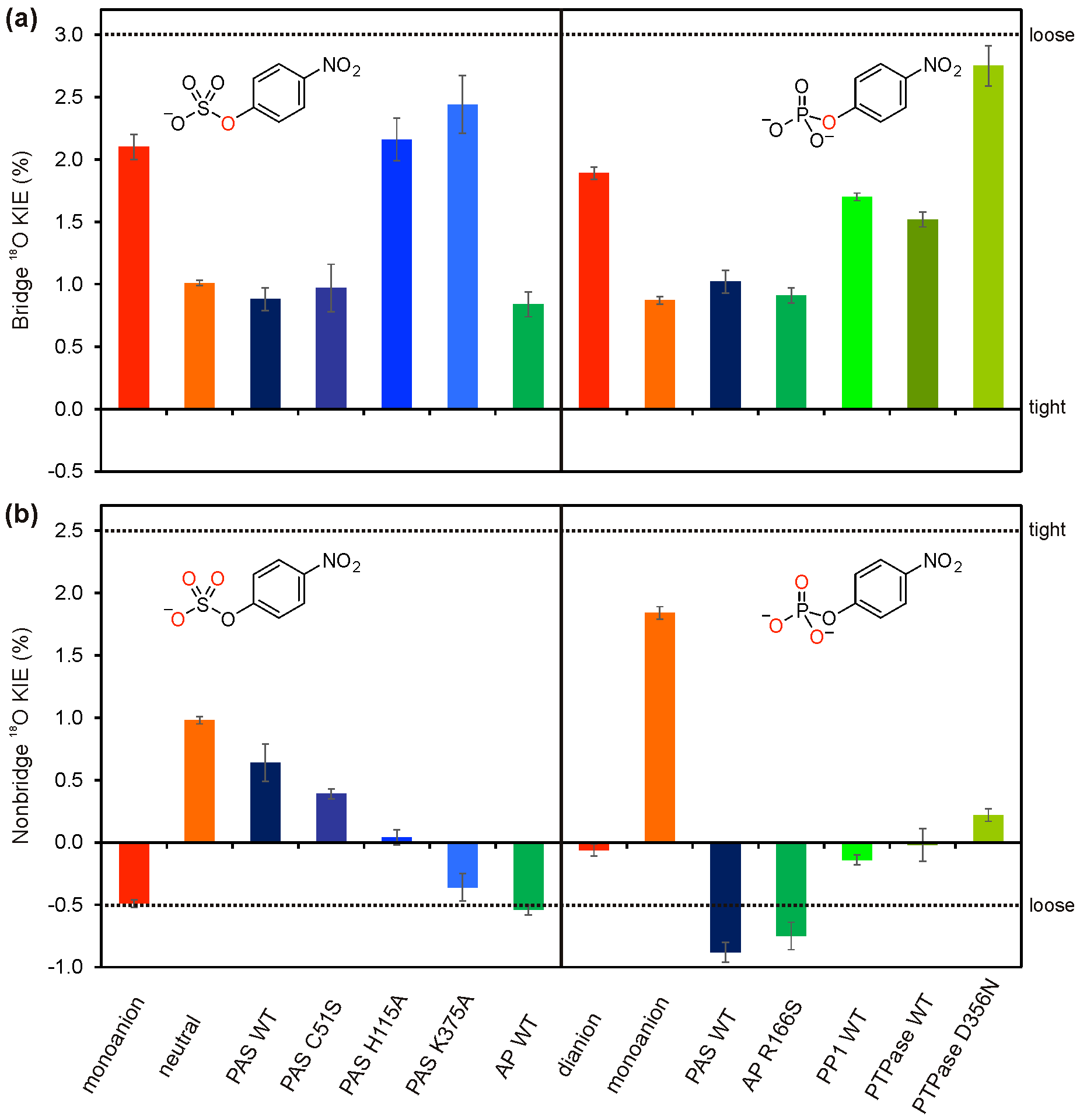
Kinetic Isotope Effects (KIEs) for PAS-catalyzed hydrolysis of sulfate monoester **1d** and phosphate monoester **2d**. The values are listed in Table 3 (PAS and solution data) and Table S15 (all other enzymatic data, SI). (**a**) The ^18^O-KIEs for the bridging (or leaving group) oxygen. (**b**) The ^18^O-KIEs for the nonbridging oxygens. The dotted lines indicate extremes of ^18^O KIEs for tight (^18^O bridge = 0%, ^18^O non-bridge = +2.5%) and loose (^18^O bridge = +3.0%, ^18^O non-bridge = −0.5%) transition states.^90^ AP: *E. coli* Alkaline phosphatase,^20^ PP1: Protein Phosphatase 1,^57^ PTPase: *Yersinia* Protein Tyrosine Phosphatase.^53^

In the PAS WT-catalyzed hydrolysis of phosphate monoester **2d**, the ^15^(*V*/*K*) and ^18^(*V*/*K*)_bridge_ KIEs also indicate near complete neutralization of the charge on the leaving group, and significant P-O_bridge_ bond fission (Table 3, Figure 4a). The ^18^(*V*/*K*)_nonbridge_ KIE for PAS WT-catalyzed hydrolysis of phosphate monoester **2d** was more inverse than for the uncatalyzed solution reaction of the dianion (0.9912 *vs* 0.9994; Table 3, Figure 4b), and is similar to the value for its hydrolysis by the superfamily member AP R166S (0.9925).^35^ In the case of AP the inverse shift in this KIE was attributed to interactions of the phosphoryl group with the metal ions and hydrogen bonding residues at the active site. In the PAS reaction similar interactions are possible (Figure 1). For the PAS WT-catalyzed hydrolysis of sulfate monoester **1d** the ^18^(*V*/*K*)_nonbridge_ is normal (1.0064, or + 0.64%, Table 3 and Figure 4b), in contrast to the inverse ^18^(*V*/*K*)_nonbridge_ for the uncatalyzed hydrolysis of the 1d monoanion^92^ (−0.49%). This value is closer to the ^18^(*V*/*K*)_nonbridge_ for the hydrolysis of the neutral sulfate monoester^92^ (+0.98%). However, in this case, the normal KIE arises from deprotonation of the sulfuryl group (i.e. proton transfer from S-O-H to the leaving group). In previous investigations of the TS for enzymatic sulfate and phosphate transfer the drop in the normal ^18^(*V*/*K*)_bridge_ was accompanied by a more inverse^20^ or mostly unchanged ^18^(*V*/*K*)_nonbridge_ KIE^50,51,53–55,57,93^ (compared to the ^18^(*V*/*K*)_nonbridge_ for uncatalyzed phosphate dianion hydrolysis, i.e. ≤0.9994), and suggests that the TS in these enzymatic reactions is still largely dissociative despite the less negative β_leaving group_^obs^ and drop in normal ^18^(*V*/*K*)_bridge_ KIE compared to the uncatalyzed reaction, which arise from neutralization of the charge developing on the leaving group. The normal ^18^(*V*/*K*)_nonbridge_ is unlikely to arise from the same origin as in the uncatalyzed hydrolysis of the neutral sulfate monoester, since protonation of the sulfuryl group oxygens will not occur, except under extremely acidic conditions. A more plausible explanation is that the PAS WT-catalyzed TS is more associative than the solution reaction of the monoanion. The latter scenario could also partly explain the unexpectedly large effect of the removal of residues that only interact indirectly with the non-bridging leaving group oxygen on β_leaving group_^obs^.

The PAS K375A mutation results in a reaction for which the ^15^(*V*/*K*) and ^18^(*V*/*K*)_bridge_ KIEs are largely identical to those of the uncatalyzed hydrolysis (Table 3, Figure 4b), suggesting this residue is largely responsible for leaving group neutralization in the TS. This is consistent with the effect of the mutation on the β_leaving group_^obs^, and similar to what has been observed for the removal of an aspartic acid that performs a similar role in PTPs.^51,53,54^ The data suggest that this residue either protonates the leaving group directly, or, that its mutation results in a dislocation of H211 (Figure 1), interfering with its function in this role, implying synergy between these two residues, as discussed earlier. The ^18^(*V*/*K*)_nonbridge_ for PAS K375A is identical to the uncatalyzed hydrolysis, which could be explained by the loss of coordination of K375 to the sulfuryl group.

The relatively modest effect of the C51S mutation on ^15^(*V*/*K*) and ^18^(*V*/*K*)_bridge_ compared to the wild type reaction is consistent with the modest effect of this mutation on β_leaving group_^obs^. As stated above, the TS of the PAS WT-catalyzed sulfate monoester hydrolysis has a more associative character than in the uncatalyzed reaction, i.e. the nature of the nucleophile is thought to be more important, based on the change to a normal ^18^(*V*/*K*)_nonbridge_. However, the change in nucleophile from fGly to serine has a much smaller effect on ^18^(*V*/*K*)_nonbridge_, than the removal of K375, which is thought to interact directly with the nonbridging oxygens (Figure 1).

The mutation of H115 to alanine has a large effect on the KIEs compared to WT, despite the absence of direct TS interaction between H115 and the substrate (Table 3, Figure 4). The ^18^(*V*/*K*)_bridge_ is nearly as large as that observed for PAS K375A. The ^15^(*V*/*K*) shows partial charge neutralization on the leaving group, intermediate between the WT reaction and that of K375A. A possible explanation is suboptimal orientation of the substrate relative to the residues mainly responsible for leaving group stabilization, K375 and H211. The reaction of the H115A mutant shows a ^18^(*V*/*K*)_nonbridge_ that is also intermediate between the WT and K375A, also consistent with less than optimal coordination of the non-bridging oxygens to K375. The fact that mutation H115A has a larger effect on ^18^(*V*/*K*)_nonbridge_ than C51S confirms that the large change in ^18^(*V*/*K*)_nonbridge_ for the PAS WT-catalyzed reaction compared to the sulfate monoester monoanion uncatalyzed hydrolysis is most likely not dependent on the nucleophile, but the result of interactions between the nonbridging oxygens and positively charged functional groups (K113, K375 and Ca^2+^). The nature of the contribution of the latter interactions to TS stabilization appears to be unique for PAS WT-catalyzed sulfate monoester hydrolysis, since the effect of the enzyme on the ^18^(*V*/*K*)_nonbridge_ for phosphate monoester hydrolysis is completely different. However, both reactions show a similar degree of leaving group stabilization.

### Leaving group dependence in D_2_O

The considerably less negative β_leaving group_^obs^ and a ^15^N KIE near unity for the PAS WT-catalyzed sulfate monoester hydrolysis could be caused by protonation of the leaving group oxygen, as in pTPs.^51,53–55,58^ Lewis acid charge neutralization by metal ions can have the same effect.^20^ The available data for PAS all point to K375 as the main residue responsible for stabilization of charge that develops on the leaving group oxygen in the TS. Direct proton transfer involving lysine as the donor in enzyme active sites is rare, and charge compensation by Lewis acid interaction with the cationic charge of the protein shared between H211 and K375 is a potential alternative. The PAS WT-catalyzed sulfate monoester hydrolysis in D_2_O is more sensitive to the leaving group (β_leaving group_ is more negative) compared to the same reaction in H_2_O (Figure S12). The observed difference in leaving group dependence (Δβ_leaving group_^H-D^) is +0.06±0.03, which corresponds to a *k*_H_/*k*_D_ ratio ranging from ~1.7 at p*K*_a_^leaving group^ = 5.5 to ~3.3 at p*K*_a_^leaving group^ = 10. These data suggest that with increasing demand for leaving group stabilization, a proton transfer event becomes more rate-limiting. This would suggest that the degree of S-O_bridge_ bond fission during the TS is reduced with increasing leaving group ability (i.e. S-O_bridge_ bond fission is almost complete prior to proton transfer for the low p*K*_a_ leaving groups that require charge offset assistance less). However, the ^15^N KIE for PAS WT-catalyzed hydrolysis of sulfate monoester **1d** (p*K*_a_ = 7.02) suggests almost complete charge compensation on the leaving group oxygen, suggesting K375 is mainly responsible for charge compensation without transferring its proton during catalysis. For PAS K375A, there is no difference in β_leaving group_^obs^ when recorded in H_2_O or D_2_O (Figure S13). However, a p*K*_a_-independent *k*_H_/*k*_D_ of ~2 for all reactions was observed, suggesting that a leaving group independent proton transfer event is rate-limiting for this mutant. The solvent isotope effect is consistent with the suggested catalytic role for K375 as the main residue responsible for leaving group stabilization during bond-breaking in the TS, since any leaving group dependent proton transfer event will most likely be fast compared to the severely slowed down S-O_bridge_ bond fission. Indeed, the heavy-atom isotope effects suggest that these two events occur in the same step.

## IMPLICATIONS AND CONCLUSIONS

A mechanistic pathway for PAS WT-catalyzed sulfate monoester hydrolysis (Figure 1b) had been suggested by structural analysis, but can now be firmly established on the basis of the results presented in this study, including a reassessment of the contributions of the active site residues. For example, leaving group stabilization by the proposed general acid H211 is much less important than that of K375 (based on the much smaller effect on β_leaving group_^obs^ upon alanine scanning). The combined effect of removing both these residues was larger than the sum of its effects in wild-type enzyme, suggesting a high degree of interdependence between these two residues with regard to leaving group stabilization, possibly by sharing a proton (Figure 5). The LFERs for *k*_cat_ and 1/*K*_M_ for PAS WT-catalyzed sulfate monoester hydrolysis and the pre-steady-state measurements with sulfate monoester **1d** suggest that the leaving group dependent sulfate ester bond fission (S-O_bridge_ fission; step 1 in Figure 1b) is much faster than the cleavage of the hemiacetal intermediate (step 2, Figure 1b).

**Figure 5.**
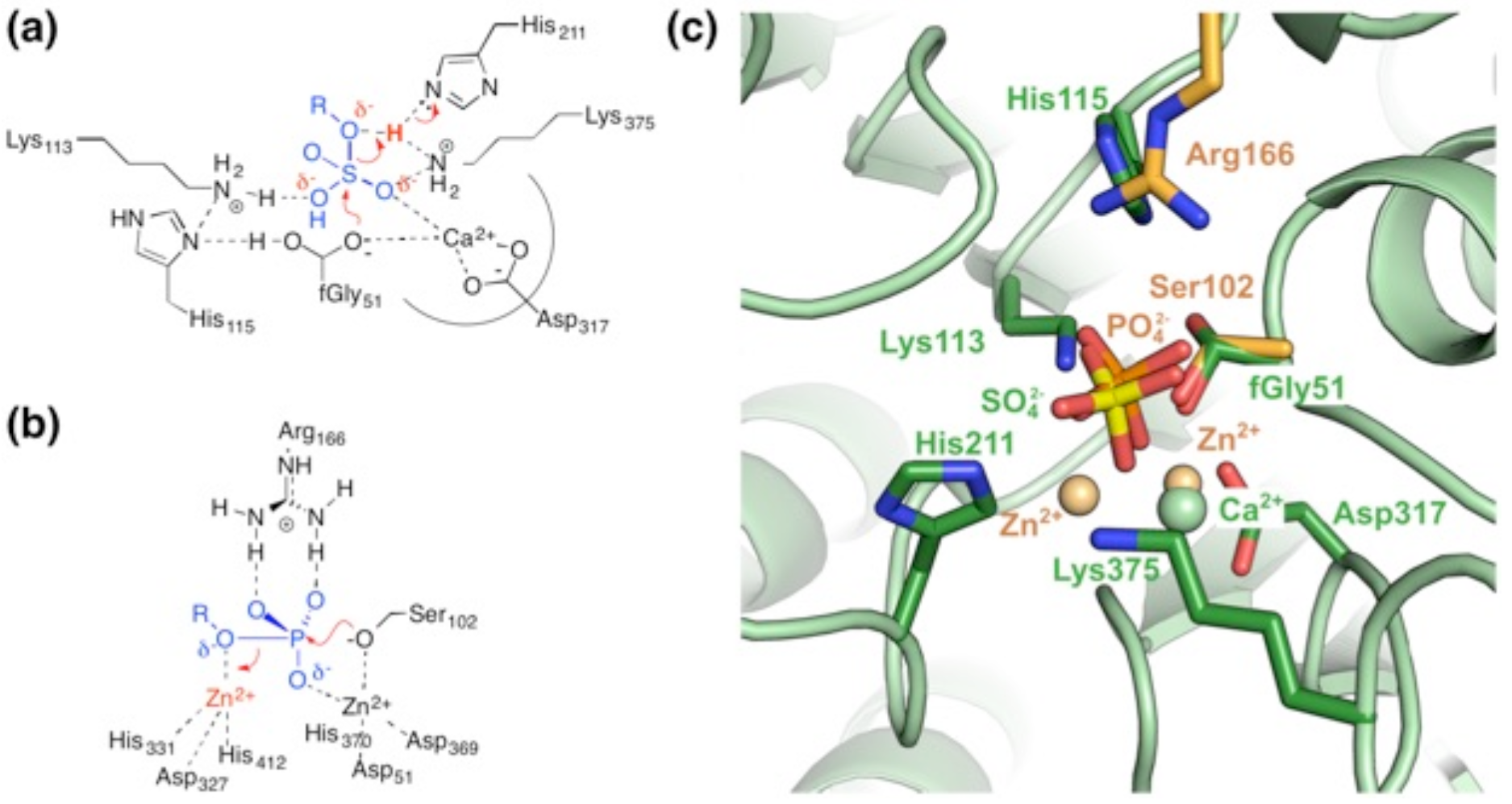
Comparison of mechanisms of leaving group stabilization in the transition states of sulfate monoester hydrolysis by PAS and phosphate monoester hydrolysis by alkaline phosphatase. (**a**) During fission of the sulfate ester bond negative charge develops on the leaving group and is offset by a proton, held by the H211-K375 pair. (**b**) In *E. coli* alkaline phosphatase (AP)^25,28^ the charge developing on the leaving group as a serine nucleophile attacks is offset by a metal ion. (**c**) Structural superposition of PAS^42^ (pdb entry 1HDH) and AP^94^ (3TG0), including bound product in the active site (inorganic sulfate and phosphate, respectively. The functionally homologous residues align well, showing that the second divalent metal ion (Zn^2+^) of AP is providing charge offset in a similar position as the proton held by H211 and K375 in PAS.

The strongest effect on leaving group dependent catalysis was seen for the K375A mutants (Δβ_leaving group_^obs^ = +0.61, Figure 2f, Table 2), further underlining that K375 is the most important residue for stabilization of the negative charge that develops on the leaving group oxygen during catalysis. This conclusion was also supported by the following observations: (*i*) The ^15^N and ^18^O_bridge_ KIEs for mutant K375A are essentially the same as for the solution reaction. The wild-type showed almost complete charge compensation on the leaving group (^15^N KIE essentially unity) and a lowered normal ^18^O_brigde_ KIE (Table 3, Figure 4). (*ii*) For PAS K375A the leaving group dependent step becomes increasingly more rate-limiting with decreasing leaving group ability (Figures 2f, S10b), pointing to its involvement in leaving group stabilization. (*iii*) The unexpectedly large effects on β_leaving group_^obs^ for active site mutants C51S and H115A are dependent on the presence of K375 (Table 2), suggesting that the main function of the other active site residues is to position the substrate for optimal interaction with K375. (*iv*) Structural alignment of PAS^42^ with AP^94^ shows that K375 occupies a similar position as the divalent metal ion thought to provide leaving group stabilization in AP (Figure 5).

Compensation of charge development on the leaving group oxygen by an amino acid side chain may be expected to involve direct protonation of the leaving group oxygen. The cleavage of this S-O_bridge_ ester is dependent on, and occurs in concert with a near-complete neutralization of the charge on the leaving group for sulfate monoester **1d** (p*K*_a_ ^leaving group^ = 7.03), as evidenced by a ^15^N KIE near unity (Table 3). This would suggest near-complete proton transfer during the TS, but transfer of the proton, would only occur once the S-O_bridge_ bond cleavage is well advanced, if at all (Figures 5a and b). Charge compensation at the leaving group oxygen is near completion for PAS WT-catalyzed hydrolysis of sulfate monoester **1d**. As the proton is shared between H211 and K375 (Figures 5a and b) it may be that actual proton donation from lysine does not take place and instead transfer to H211 results in the observed compensation. Comparison of the leaving group dependences in H_2_O and D_2_O showed an increasing *k*_H_/*k*_D_ ratio for *k*_cat_/*K*_M_ with decreasing leaving group ability (Figure S12, SI), suggesting that proton transfer is becoming increasingly more important for S-O_bridge_ ester bond fission. The absence of the leaving group dependent change in *k*_H_/*k*_D_ for PAS K375A (Figure S13) further supports the importance of K375 for S-O_bridge_ bond fission, since it is in agreement with S-O_bridge_ ester bond fission being fully rate-limiting for leaving group departure in this mutant. (This solvent isotope effect is relevant for the first chemical step, while the isotope effect on *k*_cat_ reflects the overall rate-determining step, i.e. breakdown of the intermediate).

The correlation of reaction rates and the measured values for β_leaving group_^obs^ (Figure 6) establishes a direct link between charge compensation and catalytic efficiency: the more the catalytic effect of the proton is removed (as indicated by larger effective charge changes at the leaving group oxygen), the more does the overall rate suffer. This effect is more pronounced for unreactive substrates that require interaction with the proton held by K375 and H211 to a greater extent (Figure S15).

**Figure 6.**
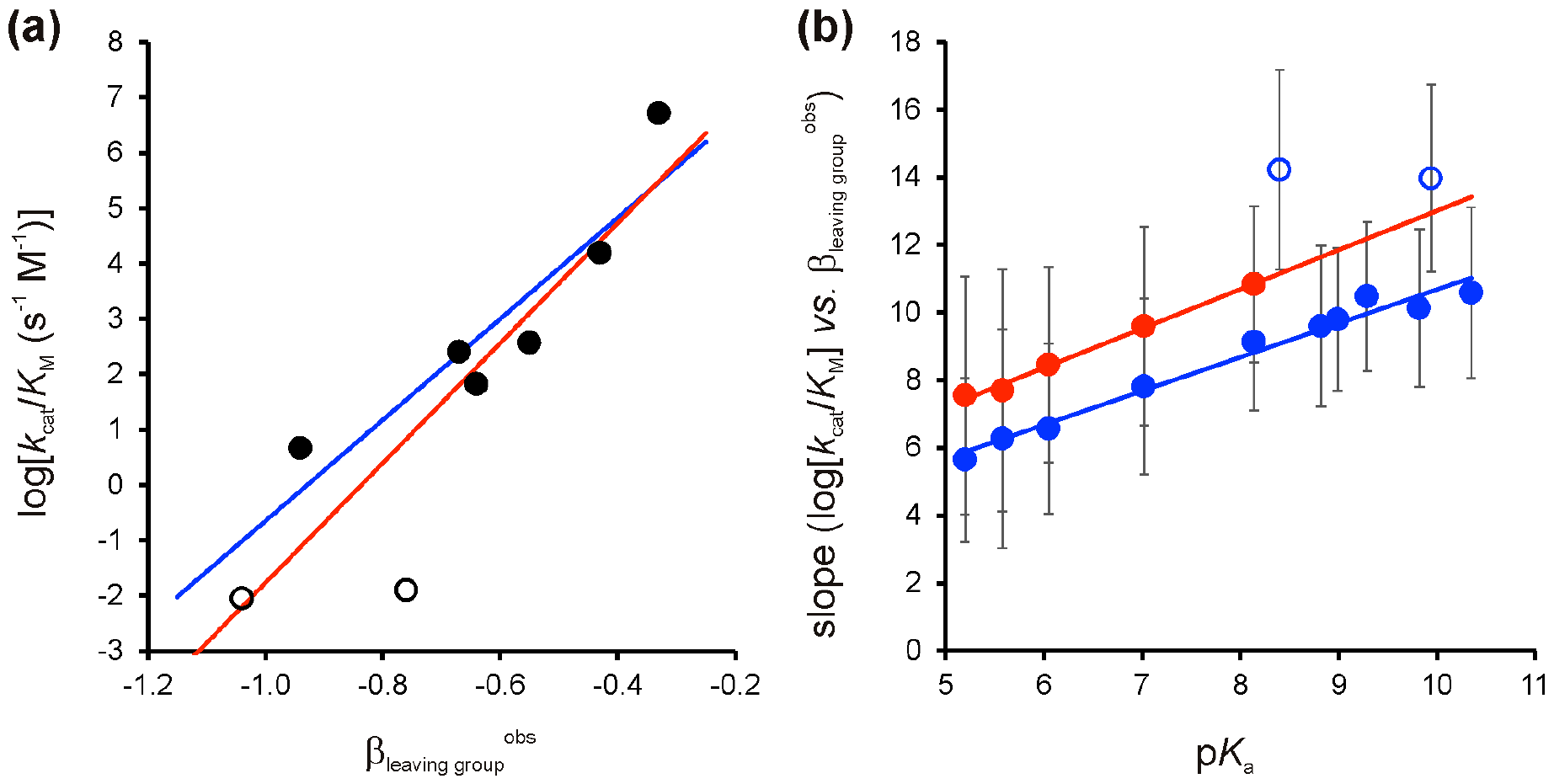
PAS variants with lower catalytic efficiencies are more sensitive to the leaving group ability. (**a**) Correlation between catalytic efficiency (represented by log[*K*_cat_/*K*_M_]) and leaving group dependency (β_leaving group_^obs^) of the various PAS variants (closed symbols: PAS WT and its single mutants, open symbols: double mutants) for 3-nitrophenyl sulfate **1e**. Linear correlations including all variants (red line, R^2^ = 0.79; p = 0.0033) and only including the single mutants and wild type (blue line, R^2^ = 0.83; p = 0.011) show slopes of 10.8±2.3 and 9.1±2.0 respectively. (b) Correlation between the response slope as shown in panel (a) and p*K*_a_ values of the substrate’s leaving group (see Figure S14 for the corresponding correlation lines). The slopes for sulfate monoesters **1f** and **1k** clearly deviate from the observed trend and were therefore not included in the fit (for more details see Figure S15, SI). Red fit: slope: 1.16±0.05; R^2^: 0.99; p: 2.2 × 10^−4^. Blue fit: slope: 1.0±0.06; R^2^: 0.97; p: <10^−4^.

An alternative mechanistic scenario in which additional steps prior to the first irreversible S-O_bridge_ ester bond fission are rate limiting for wild-type, but the chemical step becomes limiting in a mutant may be considered. Such an explanation has been advanced for FLAP endonuclease, where pH-rate profiles suggested rate-limiting physical steps after substrate binding resulted in a commitment to catalysis that suppressed the magnitude of β_leaving group_^obs^ and was reduced or eliminated by a mutation that slowed the chemistry step.^95^ However, this scenario is inconsistent with the observed KIEs on PAS catalysis. A commitment factor would reduce all the observed KIEs in equal proportion.),^89^ Thus, if making the chemical step more rate-limiting (reducing a commitment factor) were the cause of the larger Brønsted slope, the KIEs would increase in the same proportion. Instead, the K375A mutant reaction gives an ^15^N KIE that is 4-fold higher than wild-type PAS; the bridge ^18^O effect is 2.8-fold greater; and the non-bridge ^18^O effect goes from normal to inverse. The substrates in FLAP and PAS also show significant differences: DNA substrates of FLAP may require unpairing or helical arch ordering, whereas for the small molecule substrates of PAS no analogous steps are conceivable. In addition the high *K*_M_-values are consistent with reversibility and a rapid equilibrium between bound and unbound states (contrasting with the nM binding in FLAP). The LFERs and KIEs for PAS use *k*_cat_/*K*_M_ comparisons that reflect the first irreversible step of the reaction, which thus is unlikely to be associated with binding.

Although the exact residues and functional groups involved differ between several previously characterised phosphatases^20,22,23,32,35,50,51,53–55,57,58,93^ and PAS, all these enzymes rely on leaving group stabilization by charge compensation, resulting in less negative β_leaving group_^obs^ values and lowered, but still normal ^18^O_bridge_ KIEs. However, the KIEs for PAS are consistent with a more associative mechanism than previously reported phosphatases, because of the observed change of ^18^O_nonbrigde_ KIE from inverse (−0.49%) to normal (+0.62%) compared to the solution reaction (Table 3, Figure 4b). In phosphatases the small inverse ^18^O_nonbridge_ KIE was either more inverse^20^ or virtually unchanged^50,51,53,54,57,93^ (Figure 4b, Table S15). The possible change from a dissociative to a more associative TS does not apply to all PAS-catalyzed conversions, since the promiscuous enzyme-catalyzed hydrolysis of phosphate monoester **2d** showed a more inverse ^18^O_nonbridge_. Enzymatic specificity towards sulfate over phosphate monoester cannot be primarily based on overall charge, demand for leaving group stabilization or nucleophile strength, since these either provide no discrimination or will favor the more highly charged phosphate monoesters. The difference in effect on the ^18^O_nonbridge_ KIE suggest that the subtle differences in geometry between the S-O_nonbridge_ and P-O_nonbridge_ bonds are responsible for making PAS specific toward sulfate monoesters.

More generally, the methodology developed here extends the use of physical-organic approaches to enzyme catalysis further, establishing the measurement of effective charge change *via* Brønsted plots for mutants as a way assess the involvement of a particular residue in catalysis. The correlation of catalyst and substrate reactivity (Figure 6) provides a further measure of sensitivity of reactivity changes and should be a quantitative measure of the efficiency of general acid catalysis catalysis. In the future it will be interesting to contrast slopes in such double reactivity plots (as in Figure 6b) for several enzyme systems and use its slopes to quantify the sensitivity of catalytic effects in different active site arrangements.

## EXPERIMENTAL SECTION

### Sulfate monoester compounds for linear free energy relationships (LFERs) and kinetic isotope effect studies

Sulfate monoester **1d** and phosphate monoesters **2d** and **2k** (Scheme 1) were purchased from Sigma. Sulfate esters were synthesized from the respective phenol and chlorosulfonic acid, phosphate monoesters from phosphoryl chloride and the respective phenol. Detailed procedures are given in the SI. The isotopically labeled forms of 4-nitrophenyl sulfate (1d)^92^ and phosphate (2d)^52^ for measurement of kinetic isotope effects were synthesized as described previously.

### Construction of mutants

All mutants of PAS, except for mutant C51S, which was constructed previously,^1^ were made by site-directed mutagenesis according to the QuikChange protocol (Agilent), using primers listed in Table S16 (SI) and the appropriate template plasmid.

### Protein production and purification

Expression of recombinant protein from plasmid pME4322^96^ and derived mutants was done in *E. coli* BL21 (DE3) growing in LB or 2YT-medium containing 30 mg mL^−1^ kanamycin. The cells were grown at 37 °C until an OD_600_ of around 0.6-0.8 was reached. The culture was cooled to 30 (WT) or 20 °C (Mutants), IPTG was added up to 0.75 mM and the culture was grown for 4 hours at 30 °C (WT) or overnight at 20 °C (Mutants).

Cells expressing PAS were harvested by centrifugation and resuspended in 50 mM Tris-HCl pH 8.0. One tablet of Complete^®^ EDTA-Free protease inhibitor cocktail (Roche) per 12 g of wet cell pellet was added to the suspension and the cells were lysed either by using an emulsiflex-C5 homogenizer (Avestin) or by sonication. Cell-free extract (CFE) was obtained by centrifugation of the crude cell lysate at 30,000×g for 90 min. The PAS variants were purified from CFE by subsequent anion exchange (Q-sepharose), hydrophobic interaction (Phenyl sepharose), and size exclusion (Superdex 200) chromatography. All steps were performed in 50 mM Tris-HCl pH 8.0 with the appropriate additive for each step. The anion exchange chromatography was performed as described before.^1^ Protein containing fractions that eluted from the anion exchange column were pooled and the combined fractions were brought to 200 mM (NH_4_)_2_SO_4_ by adding the appropriate volume of 50 mM Tris-HCl pH 8.0 + 2 M (NH_4_)_2_SO_4_ and subsequently loaded onto a phenyl sepharose hydrophobic interaction column. The column was washed with 2 column volumes (CV) of 50 mM Tris-HCl pH 8.0 +200 mM (NH_4_)_2_SO_4_. Protein was eluted from the column with a gradient of 200-0 mM (NH_4_)_2_SO_4_ in 50 mM Tris-HCl over 5 CV followed by a further 5 CV with 50 mM Tris-HCl pH 8.0. Fractions containing active protein were pooled and concentrated into 50 mM Tris-HCl pH 8.0 to around 10-15 mg mL^−1^ protein. The concentrated protein was loaded onto a Superdex 200 prep grade gel filtration column. PAS eluted at the expected elution volume of monomeric PAS. Protein containing fractions were pooled and concentrated to 100-350 μM in 50 mM Tris-HCl and aliquoted in appropriate portions, flash frozen in liquid N_2_ and stored at −20 °C. Protein concentrations were calculated based on the molar extinction coefficient at λ = 280 nm, ε_280_ = 102790 M^−1^ cm^−1^, calculated from the amino acid sequences using ProtParam (http://expasy.org/tools/protparam.html).

### Enzyme kinetics

All data for steady state enzyme kinetics were recorded at 25 °C in 100 mM Tris-HCl pH 8.0 supplemented with 500 mM NaCl or as indicated. Observed initial rates (*V*_obs_) were determined by following an increase in absorbance at a fixed wavelength (ranging from 270 to 400 nm depending on the substrate) as a result of product formation over time in microtiterplates (Spectramax Plus, Molecular Devices) or quartz cuvettes (Varian 100 Bio). Catalytic parameters *k*_cat_, *K*_M_ and/or *k*_cat_/*K*_M_ were obtained by fitting the dependency of *V*_obs_ on substrate concentration ([S]) at a fixed enzyme concentration ([Enz]) (equation 2).

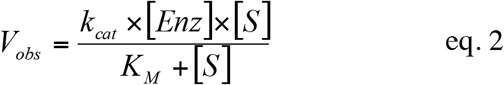

The dependency of the various kinetic parameters (*k*_cat_, 1/*K*_M_ and *k*_cat_/*K*_M_, represented by K in equation 3) on leaving group ability (as represented by their p*K*_a_’s) was fitted to equation 3 to obtain the observed Brønsted constants for leaving group dependence (β_leaving group_^obs^).

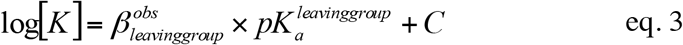

*Stopped-flow kinetics*. Fast kinetics for PAS WT-catalyzed hydrolysis of sulfate monoester **1d** were recorded using a SX20 stopped-flow setup (Applied Photophysics). No unambiguous burst phase could be detected with PAS concentrations between in 1 and 8 and 1 mM sulfate monoester **1d**, with 100 mM Tris-HCl pH 8.0, containing 500 mM NaCl at 20 °C. Data are shown in Figure S3.

### Kinetic isotope effects (KIEs)

Natural abundance **1d** or **2d** was used for measurements of ^15^(*V*/*K*). The ^18^O KIEs ^18^(*V*/*K*)_bridge_ and ^18^(*V*/*K*)_nonbridge_ were measured by the remote label method, using the nitrogen atom in *p*-nitrophenol as a reporter for isotopic fractionation in labeled bridging or nonbridging oxygen positions.^90^ The particular isotopic isomers used are shown in the SI. Isotope effect experiments used 100 μmoles of substrate, at 25 °C in 50 mM Tris buffer, pH 8.0. The substrate concentration was 19 mM and the reactions were started by addition of wild-type or mutant enzyme,1 μM for substrate **1d**, and 725 μM for substrate **1b**. After reactions reached completion levels between 40% and 60% they were stopped by titration to pH 3 with HCl. Protocols for isolation of *p*-nitrophenol, isotopic analysis, and calculation of the isotope effects were the same as previously described,^52,92^ and are described in the SI.

## ASSOCIATED CONTENT

Supporting information available. Synthetic procedures, mathematical descriptions for enzyme kinetics and expected detection limits, determination of kinetic isotope effects and data analysis. Michaelis-Menten plots for sulfate substrates hydrolyzed by PAS WT, tables of experimentally measured steady-state kinetic parameters, additional LFERs, stopped flow data and primers used for mutant construction.

## ACKNOWLEDGMENTS

This research was funded by the Biological and Biotechnological Research Council (BBSRC) and the Engineering and Physical Sciences Research Council (EPSRC) and NIH grant GM 47292 to ACH. FH is an ERC Starting Investigator, UB received a studentship from the Royal Thai Government and MG was supported by a postdoctoral Marie-Curie fellowship from the EU. We thank Marko Hyvonen for his help in the preparation of structural figures.

